# Differential Representation of Articulatory Gestures and Phonemes in Motor, Premotor, and Inferior Frontal Cortices

**DOI:** 10.1101/220723

**Authors:** Emily M. Mugler, Matthew C. Tate, Karen Livescu, Jessica W. Templer, Matthew A. Goldrick, Marc W. Slutzky

## Abstract

Speech is a critical form of human communication and is central to our daily lives. Yet, despite decades of study, an understanding of the fundamental neural control of speech production remains incomplete. Current theories model speech production as a hierarchy from sentences and phrases down to words, syllables, speech sounds (phonemes) and the movements of speech articulator muscles used to produce these sounds (articulatory gestures). Here, we investigate the cortical representation of articulatory gestures and phonemes in speech motor, premotor, and inferior frontal cortices. Our results indicate that primary motor and premotor areas represent gestures to a greater extent than phonemes, while inferior frontal cortex represents both gestures and phonemes. These findings suggest that the cortical control of speech production shares a common representation with that of other types of movement, such as arm and hand movements.

## INTRODUCTION

While the cortical control of limb and hand movements is well understood, the cortical control of speech movements is far less clear. At its most basic level, speech is produced by coordinated movements of the vocal tract (e.g., lips, tongue, velum, and larynx), but it is not certain exactly how these movements are planned. For example, during speech planning, *phonemes* are coarticulated—the *articulatory gestures* that comprise a given phoneme are modified based on neighboring phonemes in the uttered word or phrase (Whalen, 1990). While the dynamic properties of these gestures, similar to kinematics, have been extensively studied (Bocquelet et al., 2016; Bouchard et al., 2016; Carey and McGettigan, 2016; Fabre et al., 2015; Nam et al., 2010; Proctor et al., 2013; Westbury, 1990), there is no direct evidence of gestural representations in the brain.

Classically, based on lesion studies and electrical stimulation, the neural control of speech production was described as starting in the inferior frontal gyrus, with low-level, non-speech movements elicited in primary motor cortex (M1v; Broca, 1861; Penfield and Rasmussen, 1949). A more recent study of electrical stimulation sites causing speech arrest confirmed that these sites were located almost exclusively in the inferior precentral gyrus (PMv and M1v), confirming that these areas are critical for speech production (Tate et al., 2014). Recent models of speech production propose that articulatory gestures are combined to create higher-level, acoustic outputs (phonemes) (Browman and Goldstein, 1992; Guenther et al., 2006). One model (Guenther et al., 2006) hypothesized that ventral premotor cortex (PMv) and inferior frontal gyrus (IFG, part of Broca’s area) preferentially represent phonemes and that M1v preferentially represents gestures. This hypothesis is analogous to our understanding of limb motor control. Premotor and posterior parietal cortices preferentially encode for the targets of reaching movements (Hatsopoulos et al., 2004; Hocherman and Wise, 1991; Pesaran et al., 2006; Pesaran et al., 2002; Shen and Alexander, 1997), while M1 preferentially encodes reach trajectories (Georgopoulos et al., 1986; Moran and Schwartz, 1999), force (Evarts, 1968; Flint et al., 2014; Scott and Kalaska, 1997), or muscle activity (Cherian et al., 2013; Kakei et al., 1999; Morrow and Miller, 2003; Oby et al., 2013). However, the model’s hypothesized localizations of speech motor control were based on indirect evidence. The location of phonemes in PMv (Levelt, 1999) was postulated based on circumstantial evidence from behavioral studies (Ballard et al., 2000) and fMRI studies which primarily examined the syllabic — rather than phonemic — level of speech(Ghosh et al., 2008; Guenther et al., 2006; Tourville et al., 2008). This model also hypothesized that gestures are encoded in M1v based on indirect evidence of non-speech articulator movements (Fesl et al., 2003; Penfield and Roberts, 1959) and fMRI studies of syllable sequencing (Riecker et al., 2000). However, none of the modalities used in these studies had sufficient combination of temporal and spatial resolution to provide definitive information about where, and more importantly how, gestures and phonemes are encoded.

Over the last decade, electrocorticography (ECoG) has enabled identification of neural activity with high spatial and temporal resolution during speech production (Blakely et al., 2008; Bouchard et al., 2013; Cogan et al., 2014; Edwards et al., 2010; Kellis et al., 2010; Leuthardt et al., 2011; Mugler et al., 2014b; Pei et al., 2011; Roland et al., 2010). High gamma activity in ECoG in M1v concurred with Penfield’s original somatotopic mappings of the articulators (Penfield and Boldrey, 1937). Several ECoG studies have found evidence that M1v activity roughly correlates with phoneme production (Bouchard et al., 2013; Leuthardt et al., 2011; Lotte et al., 2015; Ramsey et al., 2017). Mugler et al. demonstrated that single instances of phonemes can be identified during word production using ECoG from M1v and PMv (Mugler et al., 2014b). However, the ability to decode phonemes from these areas was rather limited, which suggests that phonemes may not completely characterize the representation of these cortical areas. Some ECoG evidence exists that cortical activation differs for phonemes depending on the context of neighboring phonemes (Bouchard and Chang, 2014; Mugler et al., 2014a). Moreover, incorporating probabilistic information of neighboring phonemes improves the ability to decode phonemes from M1v (Herff et al., 2015). Therefore, these areas might demonstrate predominant representation for gestures, not phonemes. However, no direct evidence of gestural representation in the brain has yet been demonstrated.

Here, we used ECoG from M1v, PMv, and IFG to classify phonemes and gestures during spoken word production. We hypothesized that ventral motor cortex represents the movements of speech, and M1v activity accordingly predominantly represents articulatory gestures. We first examined how classification accuracy of phoneme and gestures varied with the context or position within a word. We next compared the relative performance of gesture and phoneme classification in each cortical area. Finally, we used a special case of contextual variance — *allophones*, in which the same phoneme is produced with different combinations of gestures — to highlight more distinctly the gestural vs. phonemic predominance in each area. The results indicate that gestures are the predominant fundamental unit of speech production represented in the primary motor and premotor cortical areas, while both phonemes and gestures appear to be more weakly represented in IFG, with gestures still slightly more predominant.

## RESULTS

We simultaneously recorded ECoG from M1v, PMv, and IFG (pars opercularis) and speech audio during single word, monosyllabic utterances by 9 human participants (8 with left hemispheric recordings) undergoing functional mapping during awake craniotomies for resection of brain tumors (Figure 1 and Figure S1). Lesions were remote from speech production areas and no subjects had any language or speech deficits in neuropsychological testing. We manually labeled the onset of the acoustic release of each phoneme (Mugler et al., 2014b) and we employed acoustic-articulatory inversion (AAI; Wang et al., 2014; Wang et al., 2015; see Methods) in combination with the Task Dynamic Model (Nam et al., 2012) to precisely label articulatory gesture onset. We examined z-scored activity in the high gamma (70-300 Hz) band, since this band is highly informative about limb motor (Chao et al., 2010; Crone et al., 2001; Flint et al., 2012a; Flint et al., 2012b; Mehring et al., 2004), speech (Bouchard et al., 2013; Crone et al., 2001; Pei et al., 2011; Ramsey et al., 2017), and somatosensory activity (Ray et al., 2008), and correlates with ensemble spiking activity (Ray and Maunsell, 2011) and blood oxygenation level dependent (BOLD) activity (Hermes et al., 2012; Logothetis et al., 2001).

**Figure 1.**
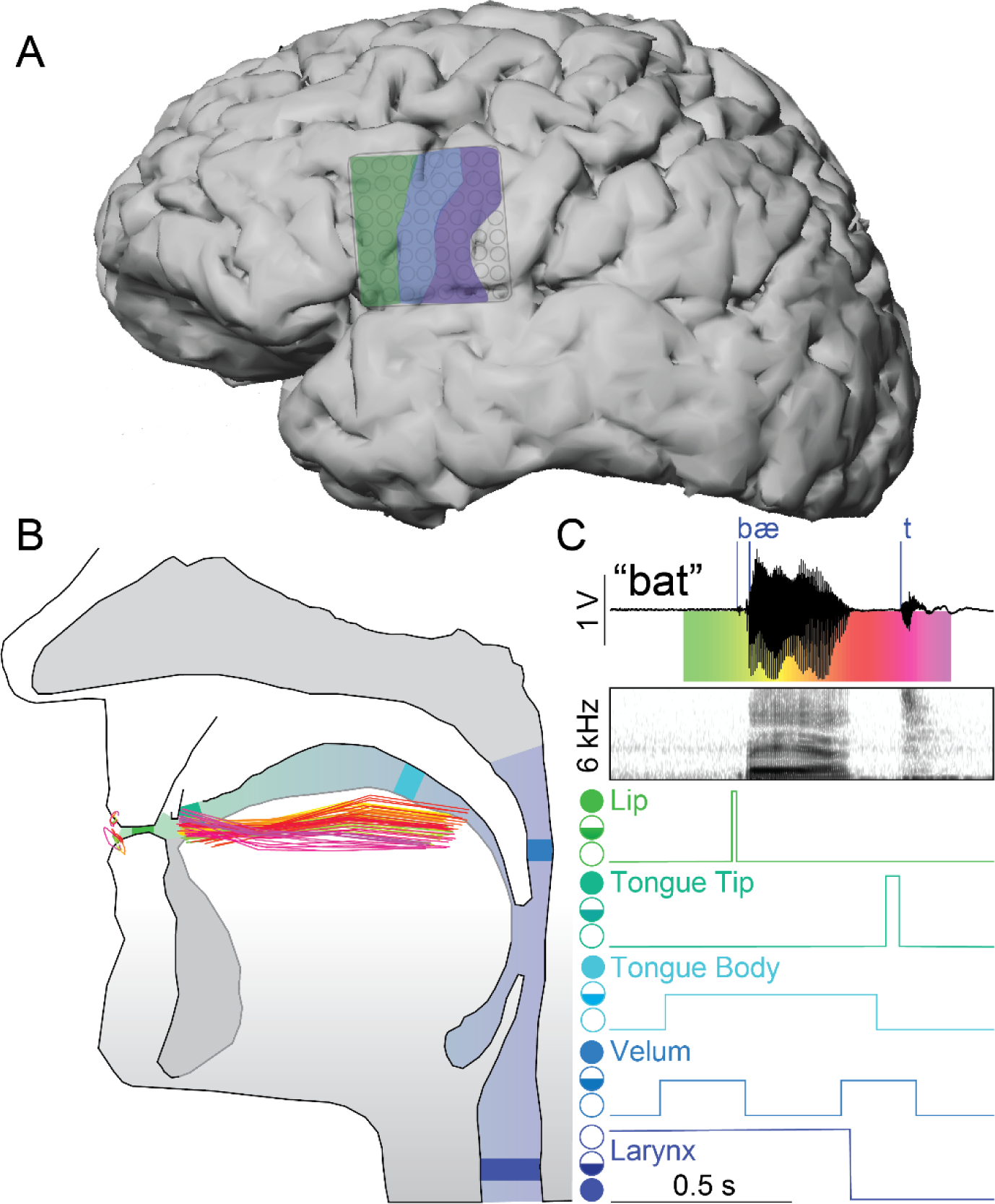
Defining phoneme and articulatory gesture onsets. (A) Cerebral cortex of participant S5 with recorded regions of speech motor cortex highlighted – IFG (green), PMv (blue), and M1v (purple). (B) Vocal tract with positions of the lips, tongue body, and tongue tip during production of a single word. Each trace represents the position, at 10-ms intervals, generated by the acoustic-articulatory inversion model, from word onset (green) to word offset (magenta; see corresponding colors in (C)). (C) Example audio signal, and corresponding audio spectrogram, from S5 with labeled phonemic event onsets (blue vertical lines) mapped to apertures along the vocal tract. Apertures for each articulator are marked from open (open circle), to critical (half-filled circle), to closed (filled circle); note that larynx has opposite open/close orientation as its default configuration is assumed to be near closure (vibrating; Browman and Goldstein, 1992).

### Phoneme-related, but not gesture-related, cortical activity varies with intra-word position

We analyzed how cortical high gamma activity varies with the context of phonemic and gestural events (i.e., coarticulation) in two subjects producing consonant-vowel-consonant words. We used the high gamma activity on each electrode to classify whether each consonant phoneme or gesture was the initial or final consonant in each word. The coarticulation of speech sounds means that phonemes are not consistently associated with one set of gestures across intra-word positions. Therefore, if gestures characterize the representational structure of a cortical area, we predicted that the cortical activity associated with a phoneme should vary across word positions. In contrast, because gestures characterize speech movements that do not vary with context, the cortical activity associated with a gesture should also be context-invariant. Therefore, we did not expect to be able to classify a gesture’s position with better than chance accuracy. We found that the high gamma activity patterns across M1v and PMv did not change with position of the gesture within a word (Figure 2A, right). In contrast, when aligned to phoneme onset, high gamma activity in M1v and PMv did vary with position within the word (Figure 2A, left). To reduce the likelihood of including cortical activity related to production of neighboring events (phonemes or gestures of lips and tongue) in our classification, we only used the high gamma activity immediately surrounding event onset (from 100 ms before to 50 ms after) to classify intraword position. Figure 2B shows an example of classification of tongue body and tongue tip closure position from all electrodes which predominantly encoded those gestures (based on single-electrode decoding of all gesture types – see Methods). Gesture classification accuracies were not larger than chance, while corresponding phonemes /k/ and /t/ were indeed larger than chance. To quantify the accuracy of classification compared to chance over electrodes, we computed the discriminability index *d′* on each electrode (Figure 2C). *d′* is the difference of means (in this case, between phoneme or gesture position and chance accuracy) divided by the pooled SD (see Methods). We computed the mean *d′* over all electrodes in M1v and PMv that were modulated with either lip or tongue movements. We found that *d′* was large for phonemes (2.3±0.6) and no different from zero for gestures (-0.06±0.6). Thus, cortical activity for gestures did not vary with context, while cortical activity for phonemes varied substantially across contexts.

**Figure 2.**
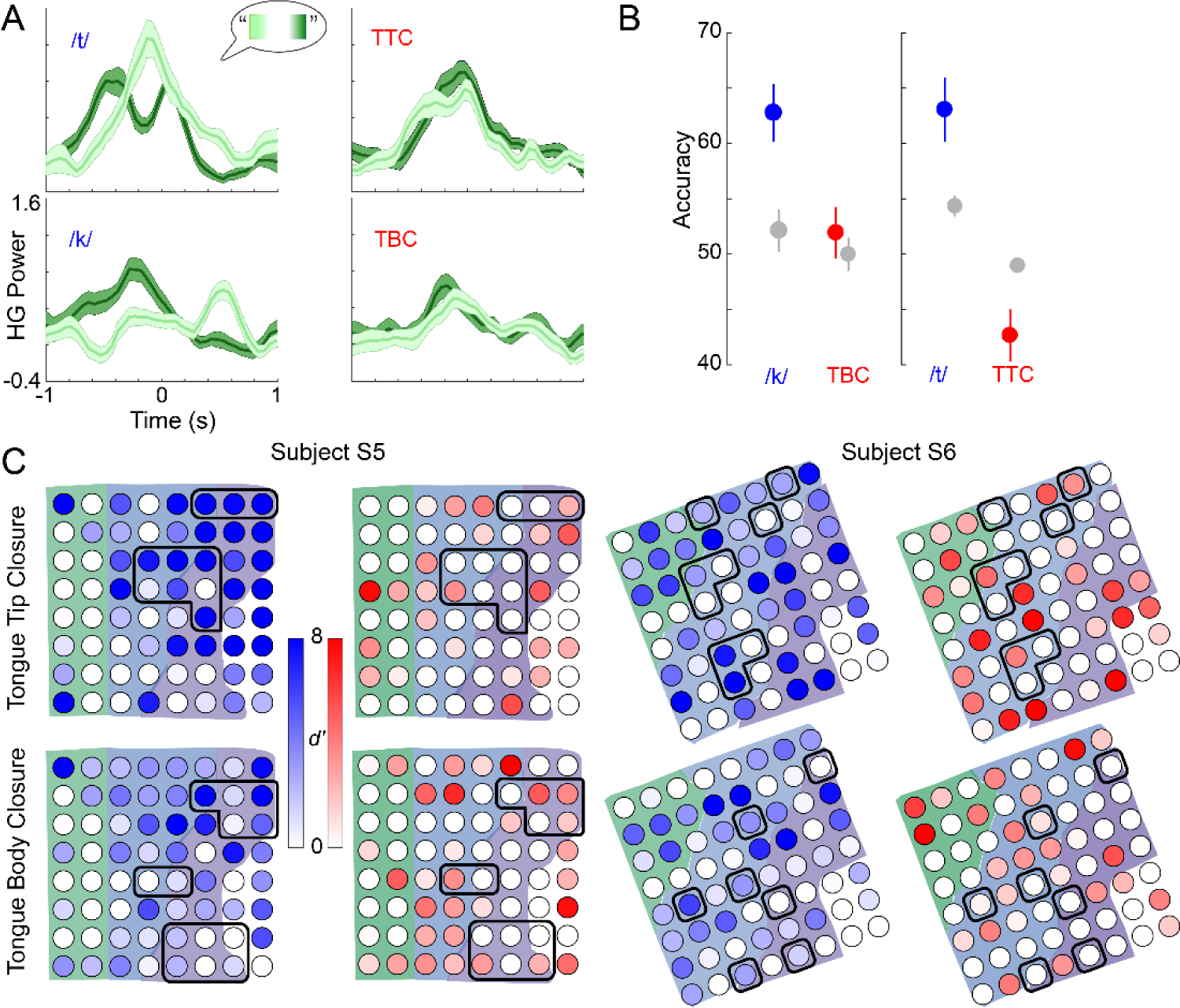
Variation of cortical activity with intraword position of phonemes and gestures. (A) Mean (±SD, shaded areas) high gamma activity on two electrodes in subject S5 aligned to onset of the phoneme (left) or gesture (right) event. Activity is separated into instances of all events (/t/ or /k/ for phonemes, tongue tip closure (TTC) or tongue body closure (TBC) for gestures) occurring either at the beginning of a word (light green) or at the end of a word (dark green). Phoneme-related activity changes with context, while gesture-related activity does not. (B) Classification accuracy (mean ± SEM) of intraword position on tongue body and tongue tip related electrodes in subject S5 for phonemes (blue), gestures (red). Gestural position classification does not outperform chance (gray), while phonemic position classification performs significantly higher than chance. (C) Cortical distribution of *d′* for differences between phonemic and gestural position accuracy and chance. Phonemic position accuracy is much higher than chance while gestural position accuracy is not on tongue tip and tongue Body related electrodes (outlined electrodes). Shaded areas correspond to cortical areas.

### M1v, PMv, and IFG more accurately represent gestures than phonemes

To further investigate sublexical representation in the cortex, we used high gamma activity from 8 participants to classify which phoneme or gesture was being uttered at each event onset. We classified consonant phonemes and gestures separately using recordings combined from motor and premotor areas (see Methods). Combined M1v/PMv activity classified gestures with significantly higher accuracy than phonemes: 63.7±3.4% vs. 41.6±2.2% (mean±SEM across subjects, p=0.01, Wilcoxon signed-rank test used for all p-values reported) as seen in Figure 3A. Gestural representation remained significantly dominant over phonemes after subtracting the chance decoding accuracy for each type (mean 34.3±3.4% vs. 17.5±2.2%, p=0.008; Figure 3B; see Methods for chance accuracy computations).

**Figure 3.**
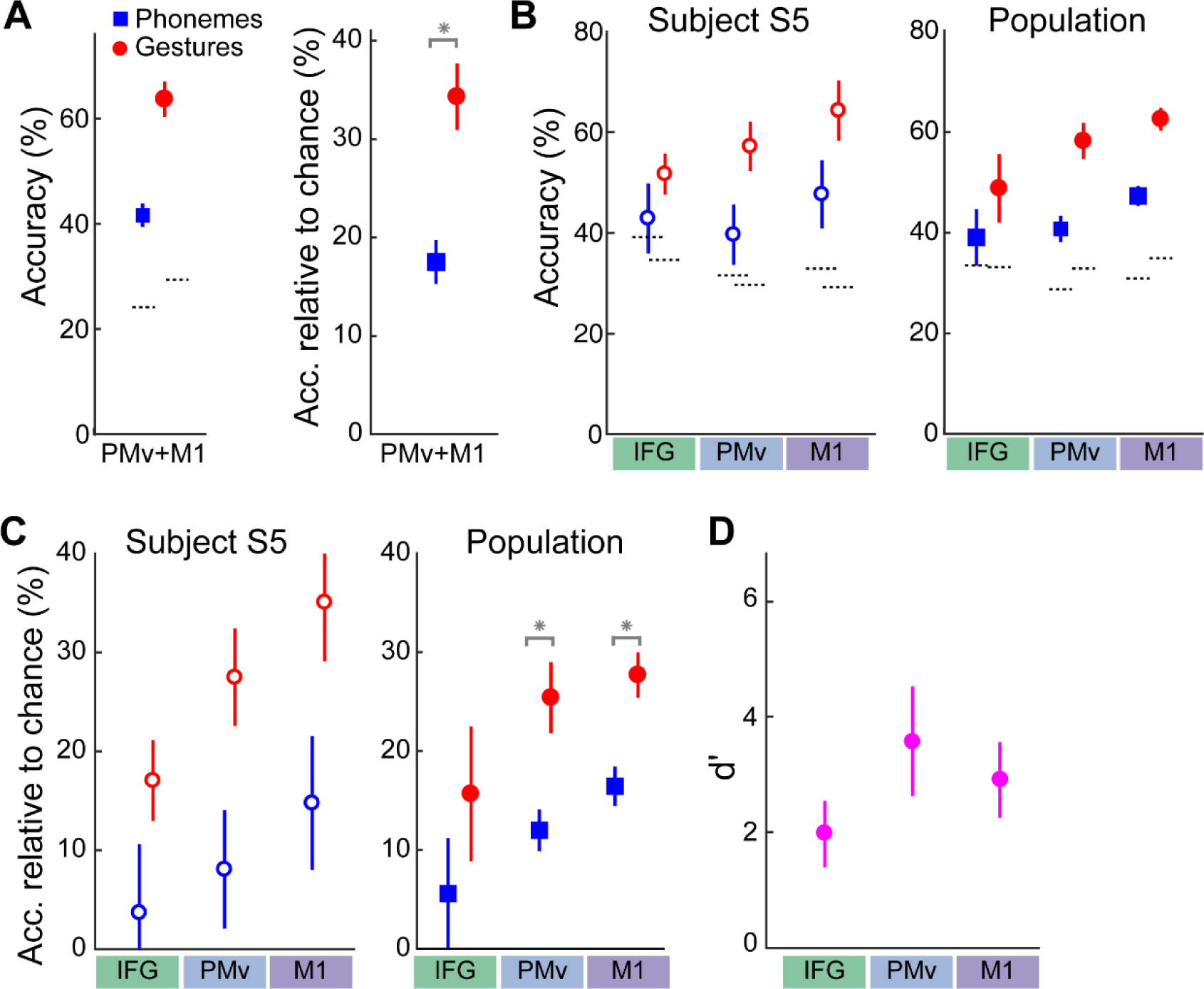
Classification of phonemes and gestures. (A) Mean (±SEM over subjects) classification accuracy using combined PMv and M1v activity of phonemes (blue squares) and gestures (red circles). Shown are both raw accuracy (left, dotted lines showing chance accuracy) and accuracy relative to chance (right). Gestures were classified significantly (*) more accurately than phonemes. (B) Classification accuracy for phonemes and gestures using activity from IFG, PMv, and M1v separately, for subject S5 (left (±SD) and population mean (±SEM, right). (C) Accuracy relative to chance in each area for subject S5 (left) and population mean (right). Gesture classification was significantly higher than phoneme classification in M1v and PMv (*). (D) *d′* values (mean±SEM over subjects) between gesture and phoneme accuracies in each area. Source data are included for A and B-D.

M1v, PMv, and IFG have been theorized to contribute differently to speech production, movements, and preparation for speech. We therefore investigated the representation of each individual area by performing gesture and phoneme classification using electrodes from each cortical area separately. Classification performance of both types increased moving from anterior to posterior areas. In each area, gestures were classified with greater accuracy than phonemes (IFG: 48.8±6.8% vs. 39.1±5.6%, p = 0.03; PMv: 58.3±3.6% vs. 40.7±2.1%, p = 0.016; M1v: 62.6±2.2% vs. 47.3±2.0%, p = 0.008; Figure 3C). This predominance remained after subtracting chance accuracy across subjects (IFG: 17.9±6.4%, p = 0.016, PMv: 25.3±12.0%, p = 0.08, M1v: 27.7±16.4%, p = 0.016; Figure 3D). The difference was significant in M1v and PMv, but not in IFG, when using Bonferroni correction for multiple comparisons. The difference in accuracy was not due to gestures having a slightly greater incidence than phonemes, as significant differences remained when we performed decoding on a dataset with maximum numbers of gesture and phoneme instances matched (data not shown). To quantify the difference further, we computed *d′* between accuracies of gestures and phonemes in each area. The *d′* values in M1v and PMv were both very high (3.6 and 2.9), while that in IFG was slightly less (2.0), suggesting decreased gestural predominance in IFG than in M1v or PMv.

### Allophone classification supports predominance of gestural representations

In four subjects, we used a specific set of spoken words from speech control literature that included allophones to amplify the distinction between phonemic and gestural representation in specific cortical areas (Buchwald and Miozzo, 2011). Allophones are different pronunciations of the same phoneme in different contexts within words, which reflect the different gestures being used to produce that phoneme (Browman and Goldstein, 1992). For example, consonant phonemes are produced differently when isolated at the beginning of a word (e.g., the /t/ in “tab”, which is aspirated, or voiceless) compared to when they are part of a cluster at the beginning of a word (e.g., the /t/ in “stab”, which is not aspirated and is acoustically more similar to a voiced /d/, Figure 4A). Using word sets with differing initial consonant allophones enabled us to dissociate more directly the production of phonemes from the production of gestures. This can be thought of as changing the mapping between groups of gestures and an allophone, analogous to limb motor control studies that used visual rotations to change the mapping between reach target and kinematics to assess cortical representation (Paz et al., 2003; Wise et al., 1998). The /t/ in “st” words was produced with high gamma activity more like a /d/ in M1 electrodes, and more like a solitary initial /t/ in PMv and IFG (Figure 4B). We trained separate classifiers for voiceless and voiced consonants (VLC and VC, respectively), and tested their performance in decoding both the corresponding isolated allophone (VLC or VC) and the corresponding consonant cluster allophone (CClA). For example, we built classifiers of /t/ (vs. all other consonants) and /d/ (vs. all other consonants) and tested them in classifying the /t/ in words starting with “st” (see Methods for details). We investigated the extent to which cluster allophones behaved more similarly to voiceless consonants or to voiced consonants. If CClAs were classified with high performance using the voiceless classifier, we would infer that phonemes were the dominant representation. If CClAs were classified with high performance using the voiced classifier, we would infer that gestures were the dominant representation (Figure 4C). If CClAs were classified with low performance by both classifiers, it would suggest that the CClA were a distinct category, produced differently from the voiced and from the voiceless allophone.

Cluster consonants behaved less like the phoneme and more like the corresponding gesture when moving from anterior to posterior in the cortex (Figure 4D and 4E). For example, in IFG and PMv, the CClAs behaved much more like the VLC phonemes than they did in M1v (p=0.6, 0.5, and 0.008 and *d′*=0.1, 0.2, and 0.4 in IFG, PMV, and M1v, respectively for performance of the VLC classifier on VLCs vs. CClAs). The CClAs behaved more like the VC phonemes in M1v than in PMv and IFG (*d′*=0.4, 0.7, and 0.3 in IFG, PMv, and M1v, respectively), although there was still some difference in M1v between CClA performance and VC performance. The CClAs were produced substantially more like VC phonemes than like VLC phonemes in M1v, which implies that M1v predominantly represents gestures. The difference between CClAs and VC phonemes suggests that the cluster allophones may represent another distinct speech sound category.

**Figure 4.**
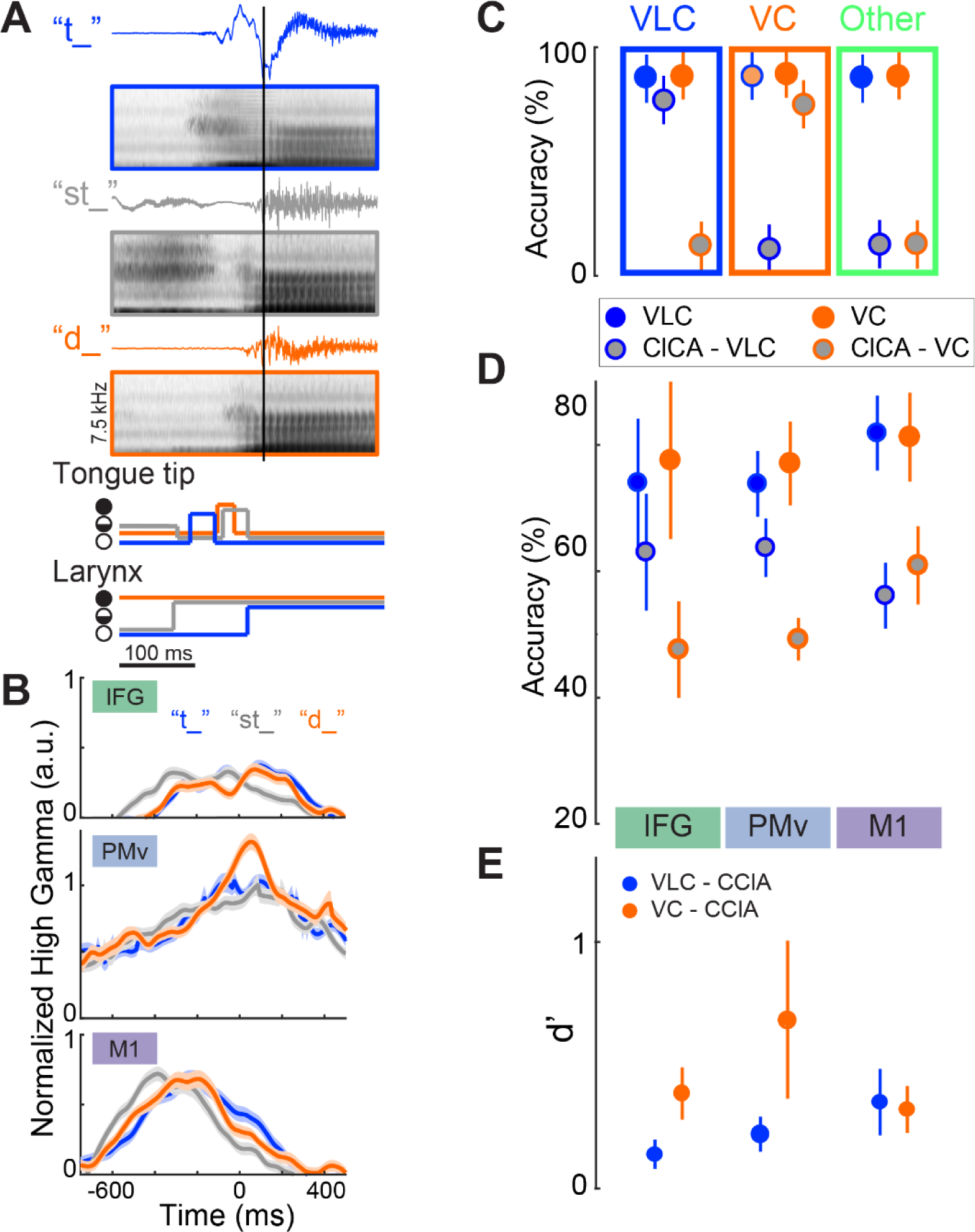
Classification of consonant allophones using ECoG from each cortical area. (A) Examples of audio waveforms, averaged spectrograms, and simplified gestural articulator trajectories for an allophone set ({/t/,/st/,/d/}) aligned to vowel onset (black vertical line). Only the trajectories for articulators that show differences for these phonemes are depicted (TT: tongue tip, Lx: larynx; filled circle: close, open circle: open, half-filled: partial closure (critical). Colors throughout the figure represent VLC (/t/, blue), VC (/d/, orange), and CClA (/st/, gray). (B) Examples of normalized high gamma activity (mean±SE) at 3 electrodes during /t/, /d/, and /st/ production in subject S5. Allophone onset is at time 0. One electrode from each cortical area is shown. CClA activity (gray) in these IFG and PMv electrodes is more similar to the VLC (blue) especially around time 0, while in M1v, it is more similar to VC (orange). (C) Schematic depicting three different idealized performance patterns in a single cortical area. Solid circles denote performance of classification of VLCs (blue) and VCs (orange) using their respective classifiers. Gray-filled circles denote CClA classification performance using the VLC (blue outline) and VC (orange outline) classifiers. High CClA performance (close to that of the respective solid color) would indicate that the allophone behaved more like the VLC or VC than like other consonants in the data set. If the CClA performed similarly to the VLC (as in the blue rectangle), it would imply that area preferentially encoded phonemes. If the CClA performed similarly to the VC (orange rectangle), the area preferentially encoded gestures. If CClA performed differently than both VLCs and VCs (green rectangle), this implied that CClAs were produced differently from either VCs and VLCs. (D) Classification performance (mean±SEM across subjects and allophone sets) in each cortical area of VLCs and CClAs in voiceless classifiers, and VCs and CClAs in voiced classifiers. CClAs show much lower performance on VLC classifiers than VLCs perform in M1v, while the performance is much closer in IFG and PMv. The opposite trend occurs with CClA performance on the VC classifiers. (E) *d′* values (mean±SEM across subjects and sets) between the singlet consonant performance and allophone consonant performance for each area; larger values are more discriminable. Blue circles: VLC vs. CClA performance using VLC classifier;s orange circles: VC vs. CClA performance using VC classifiers. In summary, CClAs perform more like VLCs and less like VCs moving from posterior to anterior. Source data are included for (D).

## DISCUSSION

We investigated the representation of articulatory gestures and phonemes in ventral motor, ventral premotor, and inferior frontal cortices during speech production. Cortical activity in these areas enabled discrimination of the intraword position of phonemes but not the position of gestures. This suggested that gestures provide a more parsimonious, and likely more accurate, description of what is encoded in these cortices. Gesture classification significantly outperformed phoneme decoding in M1v and PMv, and trended toward better performance in IFG. Cortical activity in each area, as well as in M1vand PMv combined, preferentially encoded articulatory gestures more than phonemes. Consonants in clusters behaved more similarly to the consonant that shared more similar gestures (voiced), rather than the consonant that shared the same phoneme (voiceless) in more posterior (caudal) areas, though this relationship tended to reverse in more rostral areas. Together, these results indicate that cortical activity in M1v, PMv, and possibly IFG, represents gestures to a greater extent than phonemes during production.

This is the first direct evidence of gesture encoding in the speech motor cortices. This evidence impacts theoretical models of speech production developed over decades of interdisciplinary research. The results support models incorporating gestures in speech production, such as the Task Dynamic model of inter-articulator coordination (TADA) and the Directions-Into-Velocities of Articulators (DIVA) model (Guenther et al., 2006; Hickok et al., 2011; Saltzman and Munhall, 1989). The DIVA model, in particular, hypothesizes that gestures are encoded in M1v. These results also suggest that models not incorporating gestures, instead proposing that phonemes are the immediate output from motor cortex to brainstem motor nuclei, may be incomplete (Hickok, 2012; Levelt, 1999; Levelt et al., 1999).

The phenomenon of coarticulation, i.e., that phoneme production is affected by planning and production of neighboring phonemes, has long been established using kinematic, physiologic (EMG), and acoustic methods (Denby et al., 2010; Kent, 1977; Magen, 1997; Öhman, 1966; Schultz and Wand, 2010; Whalen, 1990). Our results showing discrimination of intraword phoneme position and differences in allophone encoding confirm the existence of phoneme coarticulation in cortical activity as well. Bouchard and colleagues first demonstrated evidence of M1v representation of coarticulation during vowel production (Bouchard and Chang, 2014). Our results demonstrate cortical representation of coarticulation during consonant production. Some have suggested that coarticulation can be explained by the different gestures that are used when phonemes are in different contexts (Browman and Goldstein, 1992; Buchwald, 2014). Since gestures can be thought of as a rough estimate of articulator movements, our results demonstrating gesture encoding suggest that M1v and PMv likely encode the kinematics of articulators to a greater extent than the phonemic (or possibly acoustic) outputs.

The use of allophones enabled us to dissociate the correlation between phonemes and gestures, as a single consonant phoneme is produced differently in the different allophones. In M1v, the CClAs did not behave like either the VLC phonemes or VC phonemes, though they were more similar to VC phonemes. Overall, this suggests that the CClAs are produced differently than either VCs or VLCs, which supports previous findings. Prior to release of the laryngeal constriction, the CClAs are hypothesized to be associated with a laryngeal gesture that is absent in VC phonemes (Browman and Goldstein, 1992). Thus, it is not surprising that we observed this difference in classification between CClAs and VCs (Figure 4A). These results, therefore, still support a gestural representation in M1v as well as in PMv and IFG.

This study provides a deeper look into IFG activity during speech production. The role of IFG in speech production to date has been unclear. The classical view of Broca that IFG was involved in word generation (Broca, 1861) has been contradicted by more recent studies providing conflicting imaging evidence of phoneme production (Wise et al., 1999), syllables (Indefrey and Levelt, 2004), and syllable to phoneme sequencing and timing (Flinker et al., 2015; Gelfand and Bookheimer, 2003; Long et al., 2016; Papoutsi et al., 2009). Flinker et al. showed that IFG activity was involved in articulatory sequencing (Flinker et al., 2015). The trend toward greater accuracy in classifying gestures than phonemes using IFG activity suggests that there is at least some information in IFG related to gesture production. While our results cannot completely address IFG’s function due to somewhat limited electrode coverage (mainly pars opercularis) and experimental design, they do provide evidence for gesture representation in IFG.

These results imply that speech production cortices share a similar organization to limb-related motor cortices, despite clear differences between the neuroanatomy of articulator and limb innervation (e.g., cranial nerve compared to spinal cord innervation). In this analogy, gestures represent articulator positions at discrete times (Guenther et al., 2006), while phonemes can be considered speech targets. In arm and hand areas of M1, the reach trajectory (and arm muscle activity) is represented to a greater extent than the target of a reach (Cherian et al., 2013; Georgopoulos et al., 1986; Oby et al., 2013). This suggests that M1v predominantly represents articulator kinematics and/or muscle activity, though more detailed measurements of articulator positions (or EMG) with ECoG could demonstrate this more definitively (Bouchard et al., 2016). While we found that gesture representations predominated over phonemic representations in all 3 areas, there was progressively less predominance in PMv and IFG, which could suggest a rough hierarchy of movement-related information in the cortex (although phonemic representations can also be distributed throughout the cortex (Cogan et al., 2014)). We also found evidence for encoding of gestures and phonemes in both dominant and non-dominant hemispheres, which corroborates prior evidence of bilateral encoding of sublexical speech production (Bouchard et al., 2013; Cogan et al., 2014). This analogous organization suggests that observations from studies of limb motor control may be extrapolated to other parts of motor and premotor cortices.

Brain machine interfaces (BMIs) could substantially improve the quality of life of individuals who are completely paralyzed, or “locked-in,” from neurological disorders such as amyotrophic lateral sclerosis, brainstem stroke, or cerebral palsy. Just as the cortical control of limb movements has led to advances in motor BMIs, a better understanding of the cortical control of speech will likely improve the potential for decoding speech directly from the motor cortex. A speech BMI that could directly decode attempted speech would be more efficient than, and could dramatically increase the communication rate over the current slow and often tedious methods for this patient population (e.g., eye trackers and eye gaze communication boards, and even the most recent spelling-based BMIs (Brumberg et al., 2010; Chen et al., 2015; Pandarinath et al., 2017)). While we can use ECoG to identify words via phonemes (Mugler et al., 2014b), our results here suggest that gestural decoding would outperform phoneme decoding in BMIs using signals from M1v and PMv. The decoding techniques used here would require modification for practical “closed-loop” implementation, though simple repeatable signatures related to phoneme production have already been shown to be useful for real-time control of simple speech sound-based BMIs (Brumberg et al., 2013; Leuthardt et al., 2011). Improving our understanding of the cortical control of articulatory movements moves us closer to a viable cortical speech interface that can decode intended speech movements in real-time.

A more accurate understanding of the cortical encoding of sublexical speech production could also improve identification of functional speech motor areas. More rapid and/or accurate identification of these areas using ECoG could help to make surgeries for epilepsy or brain tumors more efficient, and possibly safer, by reducing operative time and number of stimuli and better defining areas to avoid resecting (Korostenskaja et al., 2013; Roland et al., 2010; Schalk et al., 2008). These results therefore guide future investigations into development of neurotechnology for speech communication and functional mapping.

## METHODS

### Subject Pool

Nine subjects (mean age 42, 5 female) who required intraoperative ECoG monitoring during awake craniotomies for glioma removal volunteered to participate in a research protocol during surgery. We excluded subjects with tumor-related symptoms affecting speech production, and non-native English speakers, from the study. All tumors were located at least two gyri (∼2-3 cm) away from the recording electrodes. Subjects provided informed consent for research, and the Institutional Review Board at Northwestern University approved the experimental protocols.

Electrode grid placement was determined using both anatomical landmarks and functional responses to direct cortical stimulation. Electrocortical stimulation of eloquent cortex provided *a priori* knowledge of cortex functionality and served as a “gold standard” for analysis. Areas that, when stimulated, produced reading arrest were designated as being associated with language, and areas that produced movements of the tongue and articulators were designated as functional speech motor areas. ECoG grid placement varied but consistently covered targeted areas of ventral motor cortex (M1v), premotor cortex (PMv), and inferior frontal gyrus pars opercularis (IFG). We confirmed grid location with stereotactic procedure planning, anatomical mapping software (Brainlab), and intraoperative photography (Hermes et al., 2010).

### Data Acquisition

A 64-channel, 8x8 ECoG grid (Integra, 4 mm spacing) was placed over speech motor cortex connected to a Neuroport data acquisition system (Blackrock Microsystems, Inc.). Both stimulus presentation and data acquisition were facilitated through a quad-core computer running a customized version of BCI2000 software (Schalk et al., 2004). Acoustic energy from speech was measured with a unidirectional lapel microphone (Sennheiser) placed near the patient’s mouth. Microphone signal was wirelessly transmitted directly to the recording computer (Califone), sampled at 48 kHz, and synchronized to the neural signal recording.

All ECoG signals were bandpass-filtered from 0.5-300 Hz and sampled at 2 kHz. Differential cortical recordings compared to a reference ECoG electrode were exported for analysis with an applied bandpass filter (0.53 - 300 Hz) with 75 μV sensitivity.

### Experimental Protocol

We presented words in randomized order on a screen at a rate of 1 every 2 seconds, in blocks of 4.5 minutes. Subjects were instructed to read each word aloud as soon as it appeared. Subjects were surveyed regarding accent and language history, and all subjects included here were native English speakers. All subjects completed at least 2 blocks, and up to 3 blocks.

All word sets consisted of simple words and varied depending on subject and anatomical grid coverage. Stimulus words were chosen for their simplicity, phoneme frequency, and phoneme variety. Many words in the set were selected from the Modified Rhyme Test, consisting of monosyllabic words with primarily consonant-vowel-consonant (CVC) structure (House et al., 1963). The frequency of phonemes within the MRT set roughly approximates the phonemic frequency in American English (Mines et al., 1978). The Modified Rhyme Test was then supplemented with additional CVC words to incorporate all General American English phonemes to the word set with a more uniform phoneme incidence. Consonant cluster allophone words contained initial stop consonants; each allophone example included a voiced, a voiceless, and a consonant cluster allophone word (for example, “bat”, “pat”, and “spat”; Buchwald and Miozzo, 2011).

### Signal Processing

To create features in the frequency domain, we isolated power changes in the high gamma band from the neural signal. ECoG signals were first re-referenced to a common average of all channels in the time domain. The high gamma band, most commonly used in ECoG research due to its correlation with ensemble spiking activity (Ray et al., 2008), has definitions that vary widely in the literature. We used the Hilbert transform to isolate band power in 8 linearly distributed 20-Hz wide sub-bands within the high gamma band that avoided the 60 Hz noise harmonics and averaged them to obtain the high gamma power (70-290 Hz). We then normalized and z-scored each channel’s high gamma band power changes to create frequency features for each channel.

To create features in the time domain, we segmented z-scored high gamma values for each channel from 300 ms prior to and 300 ms after onset of each event (phoneme or gesture). This created discrete, event-based trials that summarized the time-varying neural signal directly preceding and throughout production of each phoneme or gesture. Time windows for allophone feature creation were shorter (-300 ms to 100 ms) to further reduce the effect of coarticulation on the allophone classification results. The time-frequency features were then identified and sorted according to phoneme or gesture event.

### Event labeling

We used visual and auditory inspection of auditory spectral changes to manually label the onset of each phoneme in the speech signal (Matlab). To label gesture onset times, acoustic-articulatory inversion was used on the audio recordings of subjects. This technique maps articulator trajectories from acoustic data, using a model that accounts for subject- and utterance-specific differences in production. We used an articulatory inversion model, described in (Wang et al., 2015), based on a deep neural network trained on data from the University of Wisconsin X-ray Microbeam corpus (Westbury et al., 1990), with missing articulatory data filled in using the data imputation model of (Wang et al., 2014). AAI output was smoothed with a Gaussian kernel of 50 ms to reduce effects of environmental noise. Based on the target phonemes, the Task Dynamic model of inter-articulator coordination was used to generate expected laryngeal and velar movement onset times (Saltzman and Munhall, 1989). We used these onset times for each event in the speech signal to segment ECoG features.

### Event Classification and Analysis

Due to the large number of potential features and relatively low number of trials, we used classwise principal component analysis (CPCA) to reduce dimensionality of the input space and hence reduce the risk of overfitting. CPCA performs PCA on each class separately, which enables dimensionality reduction while preserving class-specific information (Das and Nenadic, 2009; Das et al., 2009). Linear discriminant analysis (LDA) was then used to determine the feature subspace with the most information about the classes. The high gamma features were then projected into this subspace and LDA was used to classify the data (Flint et al., 2012b; Slutzky et al., 2011). We used one-versus-the rest classification, in which one event class was specified, and events not in that class were combined into a “rest” group. We reported only the accuracy of classifying a given class (for example, in /p/ vs. the rest, we reported the accuracy of classifying the /p/ class, but not the “rest” class), to avoid bias due to the imbalance in “one” and “rest” class sizes. We used 10-fold cross-validation with randomly-selected test sets to compute classification performance. We repeated the 10-fold cross-validation 10 times (i.e., re-selected random test sets 10 times), for a total of 100 folds. Chance classification accuracies were determined by randomly shuffling event labels 200 times and re-classifying. We created an overall performance for each subject as a weighted average of all the events; the performance of each phoneme or gesture was weighted by the probability of that phoneme or gesture in the data set.

We limited our analysis to consonant phonemes for two reasons. First, the TADA model assumes that the larynx (or glottis) is open by default (Browman and Goldstein, 1992), which makes it very difficult if not impossible to assign meaningful onset times to this gesture, which is present in all vowels. In addition, we wished to avoid influence of coarticulation of neighboring phonemes. Therefore, we removed vowels and /s/ phonemes, as well as the glottis opening gesture, from the analysis. To ensure sufficient accuracy of our classification models, we only included phonemes or gestures with at least 15 instances, resulting in roughly the same number of phoneme classes as gesture classes (average of 15.2 phonemes and 12 gestures across subjects).

The discriminability index *d′* between two groups is defined as the difference of their means divided by their pooled standard deviation. For example, 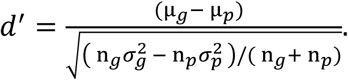 where μg is the mean of gestures, ng is the number of gesture instances minus one, and σg is the standard deviation of gesture instances, and the same symbols with subscript p stand for phonemes.

When classifying intraword position of phonemes and gestures, we examined *d′* between accuracy of phonemic or gestural position and chance accuracy. Mean values of *d′* were taken from electrodes that were related to the corresponding gesture type. This was determined by classifying among all gestures (except larynx) using the high gamma activity from each individual electrode, in 25 ms time bins, from 100 ms before to 50 ms after gesture onset. We used LDA classification (with 10x10 cross-validation repeats), since there were only 6 features for each classifier. Each electrode was denoted as related to the gesture with the highest accuracy in this classification (e.g., tongue-tip related).

## ACKNOWLEDGMENTS

We thank Robert D. Flint, Griffin Milsap, Weiran Wang, our EEG technologists, and our participants. This work was supported in part by the Doris Duke Charitable Foundation (Clinical Scientist Development Award, grant #2011039), a Northwestern Memorial Foundation Dixon Translational Research Award (including partial funding from NIH National Center for Advancing Translational Sciences, UL1TR000150 and UL1TR001422), NIH grants F32DC015708 and R01NS094748, and NSF grant #1321015.

## Supplementary Figure Legends

**Figure 1- Figure Supplement 1.** Electrode array locations for all 9 subjects. Shaded areas represent the different cortical areas: IFG (green), PMv (blue), and M1v (purple). Note that Subject 2 was implanted in the right hemisphere and so anterior-posterior direction is reversed.

**Figure 3- Figure Supplement 1.** Classification accuracy data for all subjects in Figure 3. A) Mean±SD accuracy for each subject (different symbol for each subject) for M1v and PMv electrodes combined. Chance classification performance shown as dashed line for each subject. B) Mean accuracy relative to chance for each subject for M1v and PMv electrodes combined. C and D, Same plots but using electrodes in each area for classification only.

